# The p90 Ribosomal S6 Kinases 2 and 4 promote Prostate Cancer cell proliferation in androgen-dependent and independent ways

**DOI:** 10.1101/2024.03.04.582739

**Authors:** Ryan Cronin, Aygun Azadova, Antonio Marco, Philippe P. Laissue, Greg N. Brooke, Filippo Prischi

## Abstract

Oncogenic activation of the phosphatidylinositol-3-kinase (PI3K) and mitogen-activated protein kinase (MAPK) pathways are frequent events in Prostate Cancer (PCa) that have been correlated to tumour formation, disease progression and therapeutic resistance. At the intersection of these two pathways lies the p90 ribosomal S6 kinase (RSKs) family, which regulates many proteins involved in cell survival, growth and proliferation. As such, deregulated RSKs activity has been associated with multiple cancer types, including PCa. However, the full extent of the RSKs involvement in prostate tumorigenesis remains to be determined. Here we have shown that RSKs levels are increased in PCa samples and cell lines. The RSKs were found to enhance Androgen Receptor (AR) activity, the key oncogenic driver in PCa. Indeed, all RSKs were found to interact in close proximity to the AR. However, RSK2/4, but not RSK1/3, showed changes in cell localisation following AR nuclear translocation. Consistently, silencing of RSK2/4, but not RSK1/3, inhibited PCa proliferation in an androgen-dependent and independent manner, respectively, and induced different signaling events downstream of the AR. The data suggests that RSK2 and RSK4 activity is required for PCa cell proliferation, but they are likely regulating growth via different mechanisms.

## Introduction

Prostate cancer (PCa) is predominantly a hormone-driven disease that accounts for 7% of newly diagnosed cancer worldwide (1,2). PCa growth relies on Androgen Receptor (AR) signaling, a ligand activated transcription factor that, together with the Estrogen Receptor (ER), Glucocorticoid Receptor (GR), Progesterone Receptor (PR) and Mineralocorticoid Receptor (MR), belongs to the steroid hormone receptor group of nuclear receptors (3). Upon hormone (e.g. 5α-dihydrotestosterone, DHT) binding in the cytoplasm, the AR dimerises and migrates to the nucleus where it interacts with specific DNA motifs, called Androgen Responsive Elements (AREs), present in the regulatory regions of target genes (4,5). Thus, Hormone Therapy (HT), aiming to prevent AR activation, is the backbone treatment for hormone sensitive PCa (2,6). HT is initially effective in the majority of patients, but metastatic PCa invariably develop resistance to treatments and the tumours progress to the aggressive Castrate Resistant stage (CRPC), a disease of unmet clinical need (2,4,7). Extensive studies have shown that CRPC disease progression remains mainly driven by the AR, with transcriptional activity reactivated via several possible mechanisms, including increased ligand availability, AR over-expression, expression of truncated constitutive active AR variants (AR-Vs), mutations of the AR and AR activation downstream of kinases signaling (e.g., MAPK/ERK and PI3K/Akt) pathways (4,8).

Genomic and transcriptomic data demonstrated that members of the MAPK/ERK and PI3K/Akt pathways have activating mutations or deregulated expression in 43% and 42% of primary, and 90% and 100% of metastatic PCa samples, respectively (9,10), and their concomitant activation promotes PCa progression to CRPC in human cell lines (11). At the intersection of these two oncogenic signaling pathways lies the p90 ribosomal S6 Kinase (RSKs) family, a group of four (RSK1-4) serine/threonine protein kinases that phosphorylates multiple signaling effectors involved in survival, cell cycle and transcriptional regulation (12,13). Indeed, treatment of LNCaP and PC-3 PCa cell lines with the pan-RSK inhibitor SL0101 resulted in a reduction in proliferation, which suggests that RSKs activity is required for PCa cell proliferation (14). In line with alterations in upstream pathways, RSK1 and RSK2 levels are significantly higher in human PCa samples than in normal prostate and benign prostatic hyperplasia samples (14). Further, increased RSK2 expression in LNCaP was shown to enhance the expression of prostate-specific antigen (PSA), a key diagnostic marker for PCa and an AR target gene (14). However, no direct interaction between RSK2 and AR has been reported or investigated. Instead, the enhanced regulation of PSA expression by RSK2 was shown to be dependent of the direct interaction between the kinase and the coactivator histone acetyltransferase p300 (p300 or EP300) (14).

Studies exploring the role of RSKs in PCa are scarce, and isoform specific differences in PCa disease progression are not yet defined. In fact, the four RSK isoforms perform non-redundant (sometimes opposing) cancer-specific functions (2). For example, in lung and bladder cancer cells silencing of RSK4, but not RSK1, leads to a decrease in cIAP1/2 levels, promoting apoptosis (15). Similarly, in human breast cancer (BCa) cells, downstream of IRE1α signaling, RSK2 binds and phosphorylates AMPKα2, inducing autophagy and promoting resistance to ER stress both *in vitro* and *in vivo* (16). While silencing of RSK1/3/4 has no effect on BCa cell sensitivity to ER stress (16), overexpression of RSK4 reduces invasion and metastatic potential of BCa cells by up-regulating CLDN2 and down-regulating CXCR4 (17). Here, we set out to study the crosstalk between the four RSKs and the AR in PCa. We demonstrate that expression of the four RSKs is enhanced in PCa samples. Up-regulation of the RSKs was also evident, to different extents, in a panel of cells representing different stages of the disease. Mammalian 2-hybrid assays indicated that the RSKs bind in close proximity to the AR. Importantly, we showed that RSK2/4, but not RSK1/3, co-localise with the AR in the nucleus upon androgen treatment, which also correlates with promotion of PCa cell proliferation. Overall, our findings provide insight into the specific roles of RSK isoforms in AR activation and subsequent functional outcomes.

## Methods

### Data Availability and Analysis of RSK and AR expression in patient samples

Public Prostate Cancer Patient samples (Cancer Genome Atlas, TCGA) with clinical and gene expression data were utilised for corelation and gene expression analyses. TCGA patient data sets (Tumour=498, Normal=52) were obtained from the TCGA hub at UCSC Xena (https://tcga.xenahubs.net) and cBioPortal for cancer genomics (https://www.cbioportal.org/) (18). Boxplot were generated from log2 transformed RSEM values to compare expression of AR, RSK1, RSK2, RSK3 or RK4 in 52 tumour samples with adjacent normal prostate samples. Scatter plots with Pearson’s correlation were generated from log2 transformed RSEM values to compare expression of AR with RSK1, RSK2, RSK3 or RK4 in 498 patient samples and linear models fitted to examine the strength of relationships, with a p ≤ 0.05 cut-off chosen for significance. Data analysis was performed within Rstudio version 2023.03.0+386.

### Cell lines and cells culture

The human prostate PNT1A, LNCaP, C42, C42B, 22RV1, DU145 cell lines and the COS-1 cell line were obtained from ATCC. PCa and COS-1 cells were cultured in Roswell Park Memorial Institute (RPMI) and Dulbecco’s Modified Eagle’s Medium (DMEM) medium (Invitrogen), respectively, both supplemented with 2 mm L-Glutamine, 100 units/ml penicillin, 100 mg/ml streptomycin (Sigma Aldrich) and 10% fetal bovine serum. Mibolerone (Perkin-Elmer) was resuspended in ethanol and stored at −20 °C until use. For experiments where androgen levels were manipulated, cells were seeded in phenol red free DMEM or RPMI supplemented with 2 mM L-Glutamine, 100 units/ml penicillin, 100 mg/ml streptomycin and 5% double charcoal stripped fetal bovine serum (termed stripped media).

### Plasmids

The following expression and reporter vectors have been previously described: pSV-AR, pM-LXXLL (containing SSRGL**L**WD**LL**TKDSR), pVP16-AR, PDM-LAC-Z-β-GAL, TAT-GRE-EIB-LUC, 5-GAL-LUC (19). pcDNA3.1-RSK1/2/3/4 and pM-RSK1/2/3/4 were created via amplification of each full length RSKs using primers listed in Supplementary Table S1A, and inserting into *Kpn*I/*Not*I sites of the pcDNA3.1 vector, and *Eco*RI/SalI, *Bam*HI/*Hin*dIII, *Eco*RI/*Sal*I, *Eco*RI/*Bam*HI sites of the pM vector, respectively for RSK1/2/3/4.

### Luciferase and Mammalian-2-hybrid assays

For the luciferase activity assay, COS-1 cells were co-transfected with pSV-AR and pM-LXXLL, or pSV-AR and pcDNA3.1-RSK1-4, using the calcium phosphate method (19). For the Mammalian-2-hybrid (M2H) assay COS-1 cells were co-transfected with (i) pVP16-AR and pM, (ii) pVP16 and pM-LXXLL, (iii) pVP16-AR and pM-LXXLL, (iv) pVP16-AR and pM-RSK1-4. Following transfection, cells were incubated for 20-24 hours. The media was removed, cells washed with 1 ml of PBS twice and stripped DMEM media supplemented with 1 nM Mibolerone or vehicle control (ethanol). Cells were incubated for 20-24 hours, then washed with 1 ml of PBS and lysed using 60 μl of Cell Culture Lysis Reagent (Promega). 20 μl of cell lysate was added to 20 μl of Luciferase Substrate (Promega), rotationally mixed for 15 minutes at RT and luminescence quantified on a FLUOstar Omega plate reader (BMG Labtech). For β-galactosidase, 5 μl of cell lysate was added to 50 μl of Galacton-Plus (Thermo Fisher), mixed rotationally for one hour at RT, 75 μl of accelerator was added and the reaction incubated for a further 15 minutes. Luminescence was quantified on a FLUOstar Omega plate reader. Luciferase data was normalised using β-galactosidase activity. Unless otherwise stated, experiments were carried out in triplicate and results shown are the means +/-S.E. of three or more independent experiments.

### RSK siRNA knockdown

For siRNA transfection in LNCaP cells, RNAiMAX Transfection Reagent (Thermo Fisher) was used according to manufacturers’ recommendations. siRNA with pools designed against the 3′ UTR of RSK1 (sc-29475), RSK2 (sc-36441), RSK3 (sc-36443), RSK4 (sc-39212) (Santa Cruz Biotechnology), or non-targeting control siRNA according to the manufacturer’s protocol. Briefly, the transfection with siRNA was performed when cells were 20-30% for growth assays and 70% for gene expression analysis. All siRNAs were used at a final concentration of 20 nM. Cells were supplemented with either 1 nM MIB or vehicle control 72 h post transfection. Cells were incubated for a further 24 hours for qPCR experiments or five days for WST assays.

### RNA isolation, cDNA synthesis and qPCR analysis

RNA was extracted using an Rneasy Kit (Qiagen), and reverse-transcribed with the Lunascript Reverse Transcription kit (New England Biolab) according to the manufacturer’s protocols. The quantitative real-time PCR was performed using 0.25 μM of primers (Supplementary Table S1B), 5 μL Quantitect SYBR Green PCR Kit (Thermo Fisher), and RNA/Nuclease free water. The reaction was performed at 95 °C for 2 min, 40 cycles of 95 °C for 5 s and 60 °C for 10 s using a CFX Connect Real-Time PCR System (Bio-Rad). The relative expression levels of mRNA were calculated using the 2^−ΔΔCt^ method.

### Confocal Microscopy

COS-1 cells were seeded in hormone-depleted media on coverslips in 24 well plates and left for 24 hours. Cells were transfected with 200 ng of pcDNA3.1-mCherry-RSK1/2/3/4 and/or eGFP-AR using FuGENE HD, following the manufacturer’s instructions. Cells were left for 48 hrs and treated for 1 hr +/-1 nM MIB. Cells were fixed with 2% paraformaldehyde for 10 min, washed 3 times with PBS and mounted on slides with DAPI as previously described (20). Images of COS-1 cells transgenically labelled with mCherry-RSK1/2/3/4 and eGFP-AR as well as DAPI dye to stain the nuclei were acquired using a Nikon A1R confocal with a Nikon Ti-E Eclipse microscope and equipped with a Plan Apo VC 60x DIC N2 oil immersion objective with a numerical aperture (NA) of 1.40. NIS-Elements AR software was used (version 5.21.03, build 1489) with sequential scanning of the 488 nm, 562 nm, and 639 nm laser lines and standard photomultiplier tubes for detection at 2048 × 2048 pixels resolution, variable zoom settings and 2× line averaging. Images were processed using the open-source software Fiji (21). The ratio of nucleus over cytosol in mCherry-RSK1/2/3/4 labelled COS-1 cells was determined as follows: The blue channel (Fig. S4A), revealing the nucleus stained with the DNA-intercalating dye DAPI, was used to select the nucleus using iterative minimum cross entropy thresholding against the dark background, based on (22). The selection (Fig. S4B) was stored in the region of interest (ROI) manager and applied to the magenta channel showing mCherry-RSK1/2/3/4 labelled cells. Everything outside the nuclear selection was set to black (using the ‘Clear Outside’ command), revealing only the fluorescence intensity signal within the nucleus of the magenta channel (Fig. S4C). Conversely, the nuclear selection was filled with black (using the ‘Fill’ command), leaving only the fluorescence intensity signal across the cytosol in the magenta channel (Fig. S4D). The mean fluorescence intensities of nucleus and cytosol were then measured separately and the ratio of nucleus/cytosol calculated. Thresholding results of each cell were compiled as montage and visually inspected before admitting them as datapoint. In some cases where staining was too infrequent or weak for reliable measurement, cells with robust fluorescence intensity were halved to create additional datapoints for statistical analysis. Statistical analysis of the obtained ratios of nucleus/cytosol mean fluorescence intensity was done using PlotsOfDifferences (23). Data were visualised using violin plots (with 95% compatibility intervals indicated by indentations) and a quasi-random distribution of data, along with displaying effect sizes. Corresponding p-values were produced using randomisation tests (23-25).

### Cell Proliferation Assay

Cell proliferation assay was performed in LNCaP cells as previously described (4). Cells were plated at a density of 10^4^ cells per well in 96-well plates in RPMI. After 16 h incubation, the wells were washed twice in phosphate buffered saline (PBS) before incubation for 48 h in phenol-red free RPMI supplemented with 5% charcoal stripped FBS, 2 mm l-Glutamine, 100units/ml penicillin, and 100 mg/ml streptomycin. Hormone was subsequently added and the cells incubated for a further 72 h. To measure cellular proliferation, mitochondrial dehydrogenase activity was assayed using the WST-1 reagent (Roche Applied Science Ltd, Hertfordshire, UK) as per manufacturer’s instructions. Eight wells were assayed per condition in each of three independent experiments.

### Statistical analysis

All results are from three independent experiments. ANOVA comparison test was used to analyse the differences among multiple groups. Data are presented as means ± standard deviation (SD). P values of ^*^P < 0.05, ^**^ P < 0.01, ^***^ P < 0.001, ^****^ P < 0.0001 were considered significant. All statistical data were calculated using the GraphPad Prism 8 (GraphPadSoftware Inc., La Jolla, CA, USA).

## Results

### RSKs expression levels are higher in PCa tissues

Deregulated expression of kinases has been associated with cancer initiation, promotion and progression, as well as recurrence (26). A previous study on a small cohort of PCa tissues detected higher expression of RSK1 and RSK2 levels compared to normal prostate samples (14). We sought to expand this study and investigate expression levels of all RSKs in PCa. The analysis of RNAseq data (TCGA database) revealed that the expression of all four RSK isoforms was significantly and consistently upregulated in PCa tissues compared with matched normal prostate tissues (Fig. 1A). Furthermore, in the same datasets we analysed expression levels of the AR and also found that AR expression levels were increased in PCa. This prompted us to investigate if the expression of the RSKs is correlated with AR expression. RSK2/3/4 expression was found to be positively correlated with AR expression (R = 0.6497736, 0.433637 and 0.8689225, respectively), whereas RSK1 expression was weakly negatively correlated with AR expression (R = -0.1542568) (Fig. 1B). These results suggest that that there is a connection between RSKs and AR signalling, which we set out to investigate in more detail.

**Figure 1.**
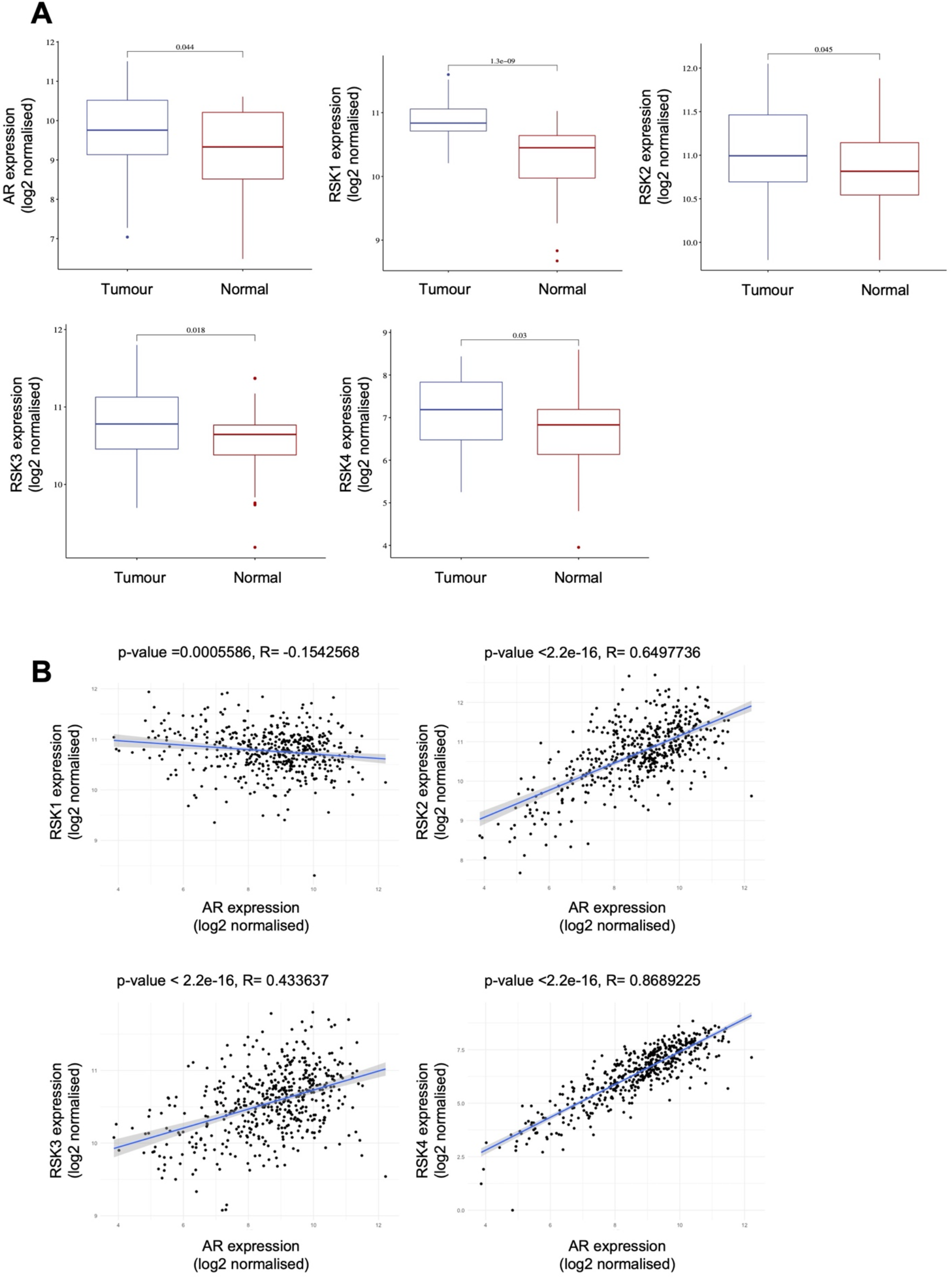
The RSKs are overexpressed in PCa. **A)** AR, RSK1/2/3/4 mRNA levels in 52 PCa samples with adjacent normal prostate samples; data were acquired from the TCGA database. The distribution of gene expression levels are shown in a box plot, where the lower and upper edges of the box represent the first and third quartiles, respectively. The horizontal line inside the box indicates the median, the whiskers represent the highest and lower datapoints and the circles represent outliers. **B**) Correlation of AR and RSK1/2/3/4 expression in 498 PCa patient samples from the TCGA database. P values < 0.05 were considered significant.

### RSKs expression is altered in PCa cell lines

RSK2 has been shown to be up-regulated in LNCaP and the growth of these cells has been shown to be partially dependent upon this kinase (14). However, little is known about the expression of the other RSKs in PCa cell lines. We therefore carried out a comprehensive study on the RSK family and investigated expression levels in a panel of PCa cell lines, representing different stages of the disease (Fig. 2). In the non-tumorigenic control cell line PNT1A, *RSK1* and *RSK2* have comparable expression levels, while *RSK3* is not expressed/barely detectable and the expression of *RSK4* is more than 14 times lower than *RSK1*. Consistent with previously published data, *RSK2* expression is enhanced (3.5 fold) in LNCaP cells, and similarly in the LNCaP derivatives C42 and C42B cells (∼2.5 fold), while its expression in 22RV1 and DU145 cells, representing more advanced stages of the disease, is not altered. *RSK2* is the most abundant isoform in all of the prostrate cell lines analysed, with the exception of 22RV1, where *RSK1* is the predominant isoform. Interestingly, RSK1 expression is similar in all lines except for 22RV1. In contrast, *RSK3* and *RSK4* are significantly overexpressed, in comparison to PNT1A, in all the PCa lines (Table S2), except for 22RV1 and DU145, respectively. Differently from what is seen in lung (13) and breast (17,27) cancers, none of the RSKs are downregulated in any of the PCa cell lines we tested. Taken together, these data highlight the presence of a complex scenario with multiple and diverse alterations in the expression of the different members of the RSK family, in line with previous observations showing isoform- and cancer-specific expression patterns and disease promoting functions (28).

**Figure 2.**
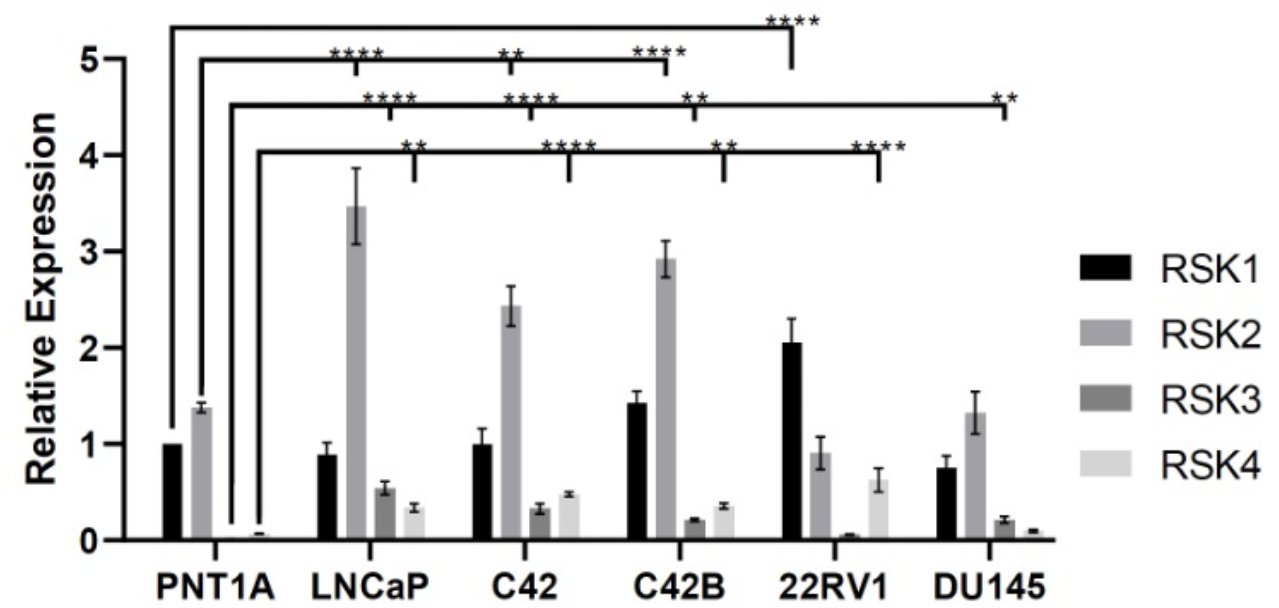
The RSKs are differentially expressed in PCa cell lines. Total RNA was isolated from a panel of PCa cell lines, converted to cDNA and RSK1/2/3/4 mRNA levels determined by qPCR. The data are shown relative to expression of RSK1 in PNT1A and are the means from 3 independent biological repeats. A schematic representation summarises prostate cell lines features and corresponding RSKs overexpression patterns. Data are presented as means ± standard deviation (SD). ANOVA was used to analyse the differences among multiple groups. ^**^P < 0.01, ^***^P < 0.001, ^****^P < 0.0001.

### RSKs regulate AR transcriptional activity

RSK1 and RSK2 have been reported to influence PCa progression by indirectly and directly altering the AR transcriptional programme, respectively. RSK2 was shown to form a complex with the co-regulator p300 and in turn activate the AR (14). Differently, RSK1 phosphorylation of YB1 induced overexpression of the AR and the constitutively active AR variant V7, key mechanisms in CRPC progression (29,30). However, no information on the interplay between AR and RSK1/3/4 is currently available (2). In order to determine the ability of the RSKs to activate the AR, we carried out transcriptional activation assays by co-transfecting each RSK into COS-1 cells (negative for the AR) with an AR expression plasmid and corresponding AR-responsive luciferase reporter construct (4). As expected, the AR was activated in the presence of the synthetic androgen mibolerone (MIB) (anon-metabolisable analogue of DHT) (Fig. 3A), confirming that the reporter system was functioning correctly in an androgen-dependent manner. RSKs did not influence AR activity in the absence of MIB, but did significantly enhance AR transcriptional activity in the presence of the ligand by approximately 6- and 3-fold, respectively for RSK1, and RSK2/3/4.

**Figure 3.**
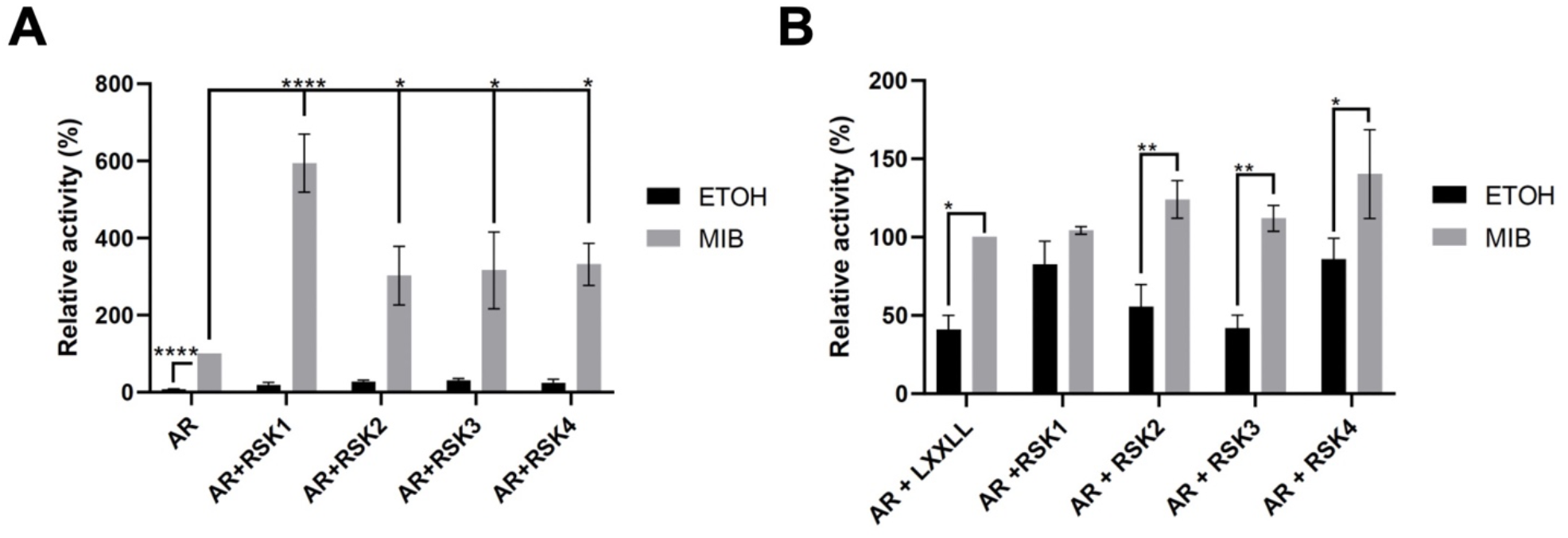
RSKs activate and interact with AR. **A**) Relative luciferase activity to show the effect of RSK1/2/3/4 upon AR activity in COS-1 cells, treated with vehicle control (ethanol, ETOH, grey bar) or mibolerone (MIB, black bar). Luciferase activity was normalised to β-galactosidase activity and made relative to AR + MIB in the absence of the RSKs. **B**) A mammalian 2-hybrid assay was performed using pVP16-AR and pM-RSK1-4. Cells were treated with vehicle control (grey bar) or MIB (black bar). Luciferase activity was made relative to pVP16-AR_+ pM-LXXLL treated with MIB. On the right, is a schematic diagram of a two-hybrid assay based on two possible AR-RSKs interaction models. Data are presented as means ± standard deviation (SD). ANOVA comparison test was used to analyse the differences among multiple groups. ^*^P < 0.05, ^**^P < 0.01, ^****^P < 0.0001.

### RSKs bind in close proximity to the AR

AR transcriptional activity can be influenced by multiple mechanisms, including post-translational modifications and regulation via co-regulators (2). Clark *et al*. (14) suggested that RSK2 regulates the AR via phosphorylation of its co-regulator p300, a mechanism which differs from that observed for the closely related ERα, where RSK2 enhances transcriptional activity via direct interaction between the kinase and the receptor (31). However, it is not known if RSKs bind directly to the AR. To investigate this and to ascertain if the RSKs enhance AR activity via formation of an activation complex, we performed mammalian 2-hybrid assays. To avoid potential false positives, due to AR transcriptional activity, the AR was used as a bait protein and fused with VP16 and the known coactivator interaction motif LxxLL (used as a positive control) and the RSKs were fused to GAL4. As expected, addition of MIB resulted in an interaction between LxxLL and the AR (Fig. 3). No signal was generated following transfection of AR-VP16 with the empty vectors, in the presence or absence of MIB, demonstrating that the system was working correctly (Fig. S4). Further, none of the RSKs, when transfected with pVP16 empty plasmid, activated the system, demonstrating that they do not have intrinsic transcriptional activity or binds to VP16. A ligand dependent interaction was identified for RSKs 2-4, with a significant increase in activity induced following the addition of MIB. In contrast, RSK1 appears to have a ligand independent interaction with the AR (Fig 3B), Taken together these data demonstrate that all of the RSKs bind in close proximity to the AR, either directly or as part of a multiprotein co-regulator complex (Fig 3B).

### The AR influences RSK2 and RSK4 cellular localisation

In an unbound state, the AR is localised in the cytoplasm where it binds chaperone and co-chaperone proteins (HSP90, HSP70, and p23) that maintains the AR in a monomeric state primed for ligand binding. Upon hormone binding, the AR dissociates from the chaperones and translocate to the nucleus (32,33). In a ligand-bound state, the AR is known to directly influence nuclear translocation of several binding partners, like PAK6 (34), β-Catenin (35), Filamin-A (36). Thus, we chose COS-1 cells (AR negative) to visualise subcellular localization of RSKs in response to AR activation. Cells expressing eGFP-AR and mCherry-RSKs were analysed by fluorescence microscopy. Confocal images show that in untreated cells, AR and RSKs are all localised in the cytoplasm and nucleus, both when expressed individually (Fig. S5) and when the AR is co-expressed in combination with each of the RSKs (Fig 5A). Upon MIB treatment, the AR migrates to the nucleus (Fig. 4A), with AR signal becoming more punctuate, which is compatible with formation of androgen-induced AR foci (37), known to be transcriptional hubs in AR positive cells (38). The AR nuclear localisation does not change in presence of the RSKs. Differently, changes in RSK2 and RSK4, but not RSK1 and RSK3, cellular localisation were observed following AR translocation to the nucleus (Fig. 4A). mChery-RSKs signal is generally more diffuse than the eGFP-AR, so we developed a new strategy to estimate significant changes for RSKs cell localisation. We determined mean fluorescence intensity in the nucleus and the cytosol separately (Fig. S4) and took their ratio as a measure of cellular localisation. RSK1 and RSK3 show little to no change in cellular localisation when expressed alone (Fig. S5) or with AR (Fig. 4B), independently of cell treatment. Differently, RSK2 shows the largest change in cell localisation upon AR ligand binding (Fig. 4B), with nearly twice as much RSK2 in the nucleus than in the cytoplasm following AR nuclear shuttling (Fig. 4C). Similarly, RSK4 shows a modest, yet significant, increase in nuclear localisation following AR nuclear translocation (Fig. 4C). Taken together these data correlate with the RSK-isoform-specific differences in AR interaction. These are likely linked to differences in affinity for the AR and/or for other RSK-isoform-specific binding partners that are part of multiprotein AR regulator complexes.

**Figure 4.**
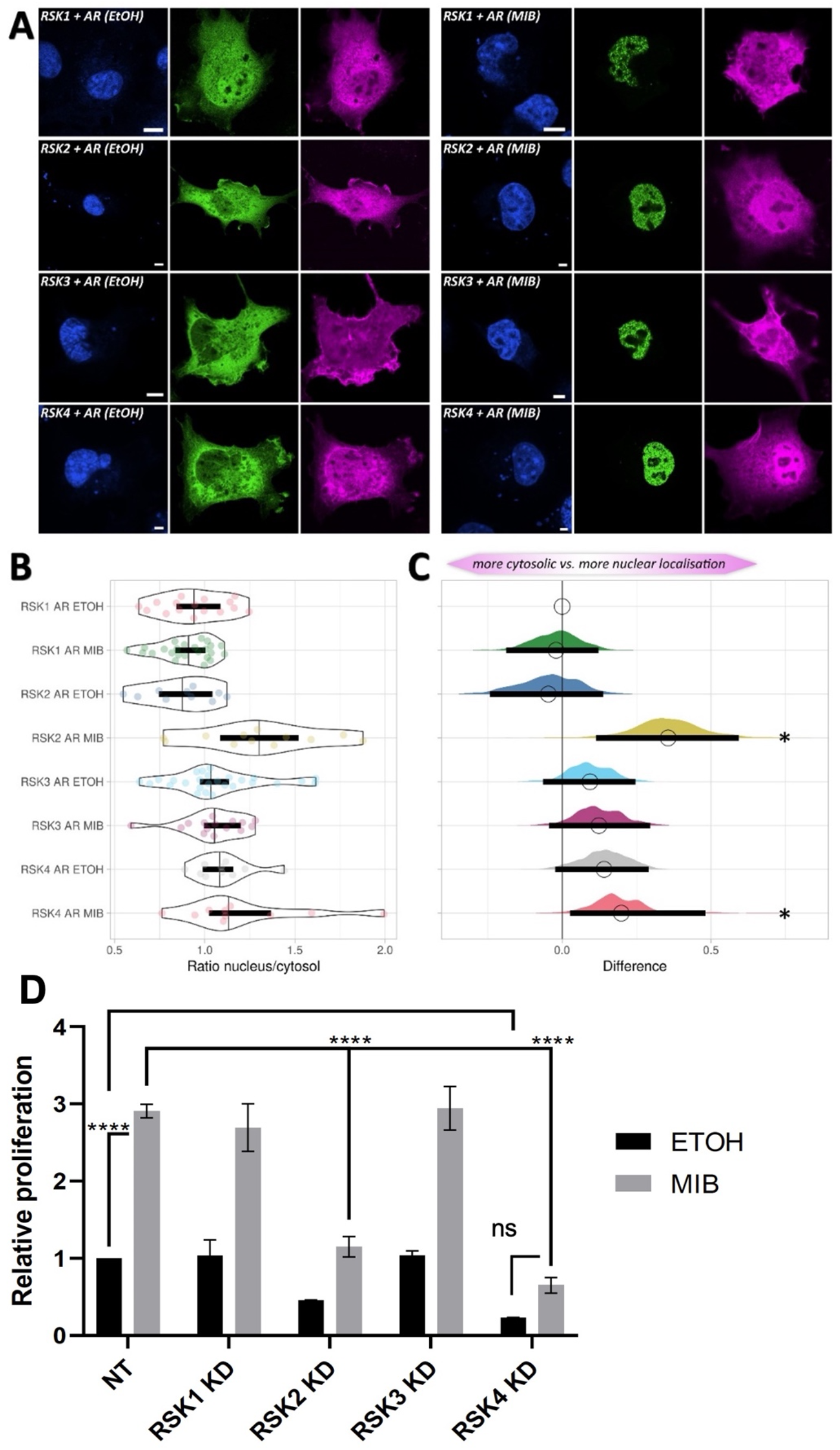
RSKs differences in cell localisation and regulation of cell proliferation. **A**) Co-expression of eGFP-AR and mCherry-RSKs. Colour channels are arranged in columns, with DAPI-stained nuclei (blue), eGFP-AR (green), and, from top to bottom, mCherry-RSK1/2/3/4 (magenta). Cells were treated with vehicle control (ETOH, left panel) or mibolerone (MIB, right panel). Scale bars indicate 10 μm. **B**) Violin plots of nucleus/cytosol ratio of mean fluorescence intensity for each of the RSKs (RSK1/2/3/4 from top to bottom), when coexpressed with eGFP-AR in cells treated with vehicle control (ETOH) or MIB. Thick black lines show the 95% compatibility interval (CI). The data points used to produce the box plots are overlaid in colour. **C**) Relative differences in the nucleus/cytosol ratio of mean fluorescence intensity for RSK1/2/3/4 (same colours as in (B)), shown as relative effect sizes. The difference between mean values is determined relative to mCherry-RSK1 co-expressed with eGFP-AR in cells treated with vehicle control (ETOH), indicated with a circle at the top. The magenta double arrow on the top of the panel indicates a stronger cytosolic localisation on the left and nucler on the right. The 95% CI is derived from the bootstrap distribution and indicated with the black vertical bars. See Table S3A-B for p-values and 95% CI. **D**) WST-1 assays were used to quantify LNCaP proliferation following RSK1/2/3/4 knock-down using siRNA. Cells were treated with vehicle control (ethanol, ETOH, grey bar) or mibolerone (MIB, black bar). Data are the means from 3 independent biological repeats. Data are presented as means ± standard deviation (SD). ANOVA comparison test was used to analyse the differences among multiple groups, with n.s., not significant. ^****^P < 0.0001.

**Figure 5.**
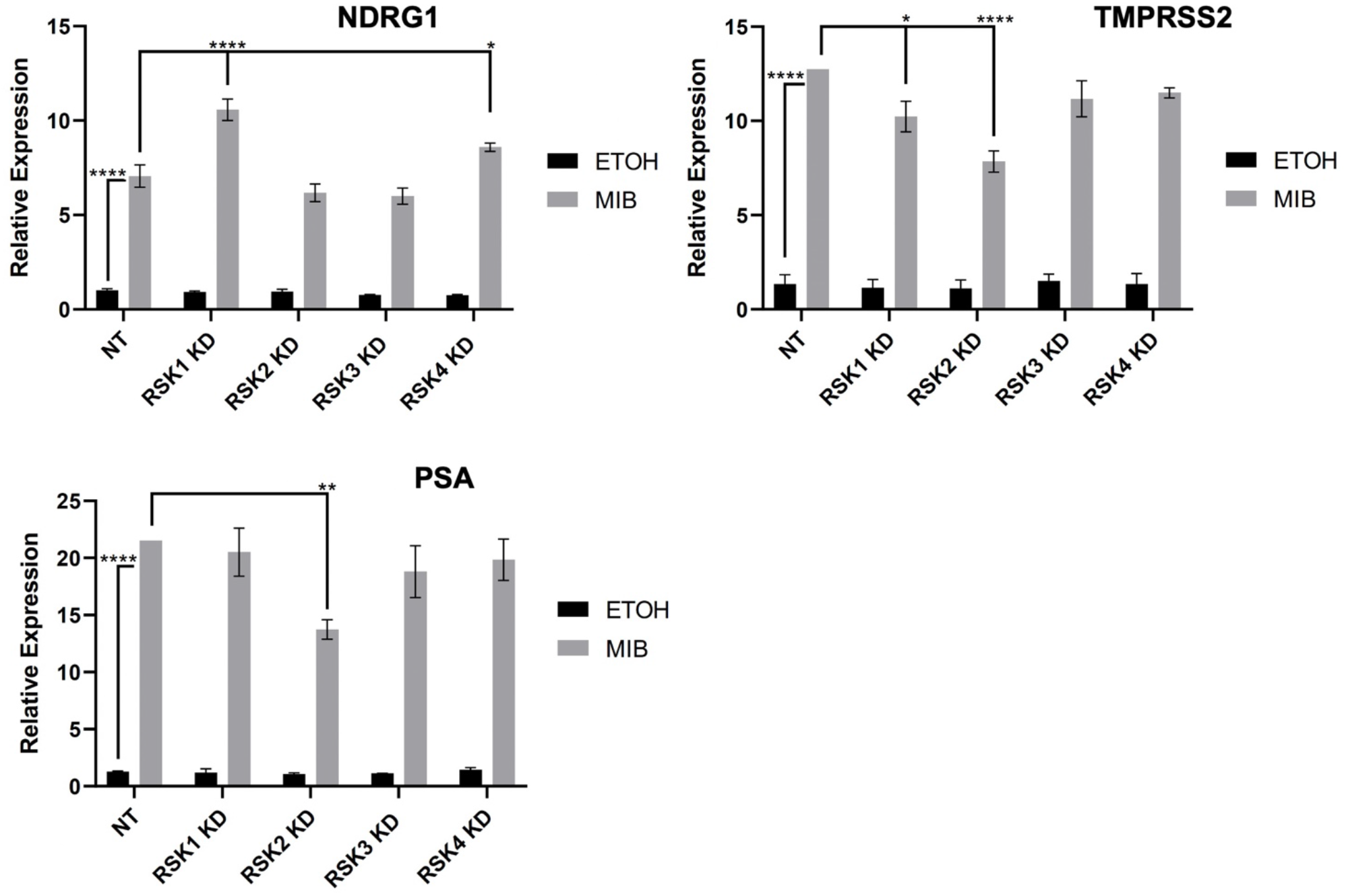
Knockdown of the RSKs has differential effects upon AR target gene expression. LNCaP cells were transfected with RSK1/2/3/4 and NT siRNAs, and treated with vehicle control (grey bar) or mibolerone (MIB, black bar). qPCR analysis of total RNA extracted from cells was used to quantify the expression of the AR target genes *NDRG1, TMPRSS2* and *PSA* (*KLK3*). Data are the means from 3 independent biological repeats ± standard deviation (SD). ANOVA comparison test was used to analyse the differences among multiple groups. ^*^P < 0.05, ^**^P < 0.01, ^****^P < 0.0001.

### RSK2 and RSK4 regulate of PCa cell proliferation

The MEK/ERK and PI3K/PDK1 pathways have been found to be over-activated in 10% and 50% of PCa patients, respectively, and they have been linked to an increase in tumour growth and disease progression (39). The RSKs are located at a major intersection point of these two signaling pathways, yet no studies have comprehensively investigated the effect of all RSK isoforms on PCa proliferation. To answer this important question, we performed siRNA knockdown of each of the RSKs in LNCaP and measured cell proliferation. MIB did not alter the expression of RSKs (Fig. S1) and successful knock-down was confirmed at the RNA level (Fig. S2). The siRNAs were confirmed to have specificity for the different RSKs, although, interestingly, knock-down of RSK2 appears to result in an up-regulation, although not-significant, of the other RSKs, suggesting a potential presence of a feedback loop.

Following knock-down, cells were treated with MIB and proliferation assays performed. As expected, a significant increase in proliferation was observed when cells were treated with MIB (Fig. 4D). Knock-down of RSK2 had no significant effect upon proliferation of non-hormone stimulated LNCaP cells, but resulted in a significant decrease in the androgen-induced proliferation of LNCaPs. Differently, silencing of RSK1 and RSK3 had no significant effect upon proliferation, in both treated and untreated cells. Interestingly, we observed a significant decrease in androgen-independent proliferation of LNCaP cells following RSK4 knock-down (Fig. 4D). Taken together these data suggest that RSK2 and RSK4, but not RSK1/3, promote LNCaP proliferation in an androgen-dependent and independent manner, respectively.

### RSKs regulation of the AR transcriptional programme

Our results, showing that the RSKs activate and bind in close proximity to the AR, albeit with some differences among the four isoforms, led us to hypothesize that these proteins contribute to the proliferation of LNCaPs by modulating the AR transcriptional program. Indeed, previous studies have shown that RSK2 increases the expression of PSA in an AR-dependent manner (14). To explore this further, we performed siRNA knockdown of each of the RSKs in LNCaP cells coupled with qPCR of known AR target genes, namely N-myc *Downstream-Regulated Gene 1 (NDRG1), Transmembrane Serine Protease 2 (TMPRSS2)* and *Prostate-Specific Antigen (PSA)* (Fig. 5). We observed key differences in target genes expression downstream of different RSK isoform signaling. In agreement with previously published data (14), silencing of RSK2 significantly reduced *PSA* expression. *TMPRSS2* expression was also reduced following RSK2 and, in smaller measure, RSK1 knockdown (Fig. 5). Both *PSA* and *TMPRSS2* are serine proteases, which promote the growth and invasion of PCa cells (40,41). Expression changes of these proteases would, at least in part, recapitulate the role of RSK2 in promoting cell proliferation. Similarly, silencing of RSK1 and RSK4 led to a significant increase in the expression of *NDRG1* (Fig. 5), a PCa metastasis suppressor (42). Alteration in *NDRG1* levels has been shown to promote PCa cell proliferation, migration, and invasion (43), which could be, in small measure, linked to the ability of RSK4 to promote cell proliferation. No changes in expression were detected for two other (*ZBT116, FKBP5*) AR target genes in response to RSKs knockdown (Fig. S6). Importantly, silencing of RSK3 had no effect on any of the AR target genes tested. This therefore demonstrates involvement in, and potentially different regulation of, AR signalling by the RSKs.

## Discussion

The RSKs are a family of serine/threonine kinases, composed of four closely related proteins (RSK1-4), that play important roles downstream of the MAPK/ERK and PI3K/PDK1 signalling cascades. RSKs activate various signalling events through selection of different substrates, many still yet to be identified, and modulate diverse cellular processes, such as cell proliferation, survival, growth and motility (44). Thus, aberrant activation of RSKs has been found to support cell transformation and tumour growth (2,28), although the mechanism of RSK action depends both on the isoform and the cancer type (13,28). Our study investigated, for the first time, the role of all four RSKs in PCa and the interplay between RSKs and the AR.

Earlier evidence from Clark et al. (14) linked elevated levels of RSK2 in LNCaP cells to PCa progression. A recent in-depth analysis of RNA-Seq data from GTEx and TCGA databases revealed that also RSK4 expression is increased in PCa (45). Using similar RNA-seq datasets, we showed that not only RSK4, but the expression of all RSKs are increased in PCa tissue when compared to healthy control tissue. Further, data from PCa cell lines representative of different stages of the disease suggests that alterations in RSKs expression may change during PCa progression, which has also been reported for other cancer types. For example, RSK1, but not RSK2/3, expression was found to increase from low to high grade gliomas (46). Similarly, RSK2 and RSK4 expression was reported to vary among different subtypes of breast cancer (47) and of renal cell carcinoma (48), respectively. While the number of comprehensive and consistent studies exploring the role of all RSKs in cancer are limited, available data suggest diverse and opposing expression patterns for the four isoforms in different tissues. In both lung and bladder cancers RSK4 has an increased mRNA expression, while RSK1 is downregulated in metastatic NSCLC (13,45,49). In breast cancer, RSK4 (45) and RSK2, but not RSK1/3 (50), expression were higher in tumour tissue than in normal breast samples. Consistently, RSK2 levels were also reported to be higher in TNBC compared with non-TNBC, and this was found to correlate with worse patient survival outcome (51). Conversely, RSK2 (52) and RSK4 (53) had a reduced expression, while RSK1 had an enhanced expression in colon cancer tissues (54). Lower levels of RSK4 in colorectal cancer also correlated with reduced survival (53). We reported a strikingly different scenario in PCa, as levels of all RSKs were found to be increased in patient samples and none of the RSKs was found to be downregulated in any of the PCa cell line analysed.

Analysis of patient datasets demonstrated that AR levels are increased in PCa, and that RSK expression correlated with AR levels. These alterations in RSK expression led us to examine the role of the RSKs in androgen signalling. Earlier studies have shown that RSK2 binds the transcriptional co-regulator p300, which in turn activates the AR transcriptional programme, resulting in a 5-fold increase in PSA expression (14). The group also demonstrated that the AR and oestrogen receptor α (ERα, another member of the steroid receptor family), are regulated differently by RSK2 (14,31). For example, RSK2 binds directly and phosphorylates the ERα on S167, enhancing ERα-mediated transcription (31). Differently, the authors excluded the possibility of a direct AR-RSK2 interaction based on the inability of RSK2 to phosphorylate *in vitro* a peptide containing residues 210-215 (formerly 203-208) of the AR. However, of the known AR phosphorylation sites (2), three lie within the RSKs consensus phosphorylation motif (R/K-x-R-x-x-S*/T*)(55): S16, S215 and S791. AKT is known to phosphorylate S215 and S791, but rather than enhance activity, phosphorylation of these sites has been shown to reduce ligand binding, destabilise the AR and reduced nuclear translocation (56). It is not known which kinase phosphorylates S16; this residue appears to play a role in AR dimerization (57). We showed that all RSKs bind in close proximity of the AR and enhance its transcriptional activity, which is compatible with both an RSK-AR direct interaction model and the formation of a multiprotein regulator complex model. Detailed interactome and/or phospho-proteomics studies in PCa are therefore warranted to better understand how the RSKs contribute to AR activation and disease progression. Indeed, studies carried out in other cancer cell types and *in vitro* have shown that RSKs have the potential to target several AR coregulators. For example, p300 can be phosphorylated also by RSK4 (13), TRIM33 by RSK1/4 (55), and STAT3, a key driver of CRPC (58), by RSK1/2 (59).

Herein, fluorescence microscopy and proliferation assays have enabled us to uncover different RSK networks in PCa cell lines. Our analysis showed that all RSKs are predominantly cytosolic and, as expected, AR translocates from the cytosol to the nucleus upon treatment with MIB, with AR signal becoming more punctate, which is compatible with formation of transcriptional hubs in the nucleus. Interestingly, RSK2 and RSK4, but not RSK1 and RSK3, showed changes in cell localisation following AR activation with MIB. An increasing number of studies suggest that RSKs subcellular distribution is regulated by binding partners (60-62), as RSKs have been found to be localised in the cytosol (45), but have also been found at the cell membrane (63) and in the nucleus (62). It is tempting to speculate that, in PCa, RSK isoform-specific binding of different AR regulators leads to differential androgen signalling, at least in part due to alterations in RSK2/4 localisation in response to AR translocation to the nucleus following ligand binding. This would also agree with the findings that RSK2/4 promotes PCa proliferation and regulates the expression of different AR target genes. RSK1 seems also able to alter indirectly the expression of AR target genes, but has little to no relevance in overall PCa cell survival. Further, RSK4 was also found to regulate PCa cell proliferation in an AR-independent manner. This is of significant relevance, as this data contributes to the growing evidence that RSK4 can promote the growth/progression of various tumours. For example, in oesophageal squamous cell carcinoma, RSK4 phosphorylates GSK-3β resulting in the activation of the β-catenin signalling pathway and promotion of cancer stem-like cell properties (64). Chrysostomou et al. (13) elegantly showed that RSK4 controls the migration and invasion of lung and bladder cancer cells, likely via phosphorylation of p300, resulting in an increase in RELA nuclear localization and enhanced NFκB activity. In addition, Pang et al. (65) found that RSK4 directly phosphorylates EZH2, activating the EZH2-STAT3 pathway and promoting resistance to inhibitors targeting EZH2 in glioblastoma. More studies are needed to better understand the role of these (or other) pathways in PCa progression and therapy resistance.

In summary, the data presented here demonstrates an important and varied role of the different RSK isoforms in AR signalling and PCa. All of the RSKs bind in close proximity to the AR and activate its transcriptional activity, but only the localisation of RSK2 and RSK4 is altered following AR activation and cause significant changes to AR target expression and overall proliferation. These findings suggest that RSK2 and RSK4 activity is required for PCa cell proliferation, but they are likely enhancing growth via different mechanisms.

### Disclosure of Potential Conflicts of Interest

No potential conflicts of interest were disclosed.

### Authors’ Contributions

**R.C**.: data curation, validation, investigation. **A.A**.: investigation. **A.M**.: formal analysis. **P.P.L**.: microscopy, formal analysis. **G.N.B**. and **F.P**.: conceptualization, supervision, project administration, investigation, writing–review and editing.

## Acknowledgments

R.C. was supported by a Peter Nicholls PhD scholarship. F.P. had funding from The Leverhulme Trust grant (RPG-2018-230). F.P. and G.B. acknowledge support from BBSRC grant (BB/X018997/1). We thank the Nikon Imaging Centre at Kings College London for help with light microscopy. The authors acknowledge the use of the High Performance Computing Facility (Ceres) and its associated support services at the University of Essex in the completion of this work. We are also thankful to Dr Kelly Coffey and Prof Luke Gaughan for critical reading of the manuscript.

---

**Table S1A.**
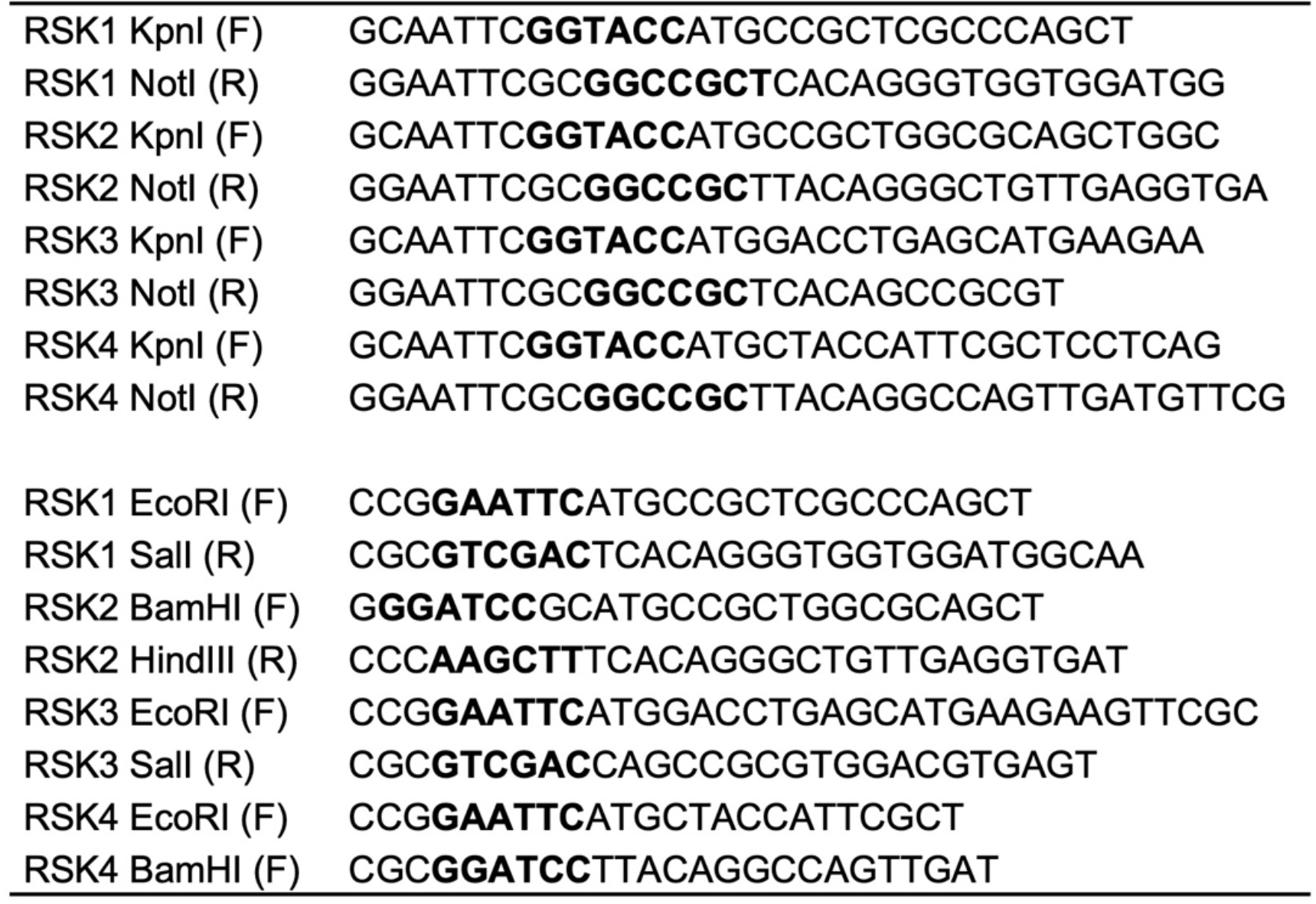
cloning primers.

**Table S1B.**
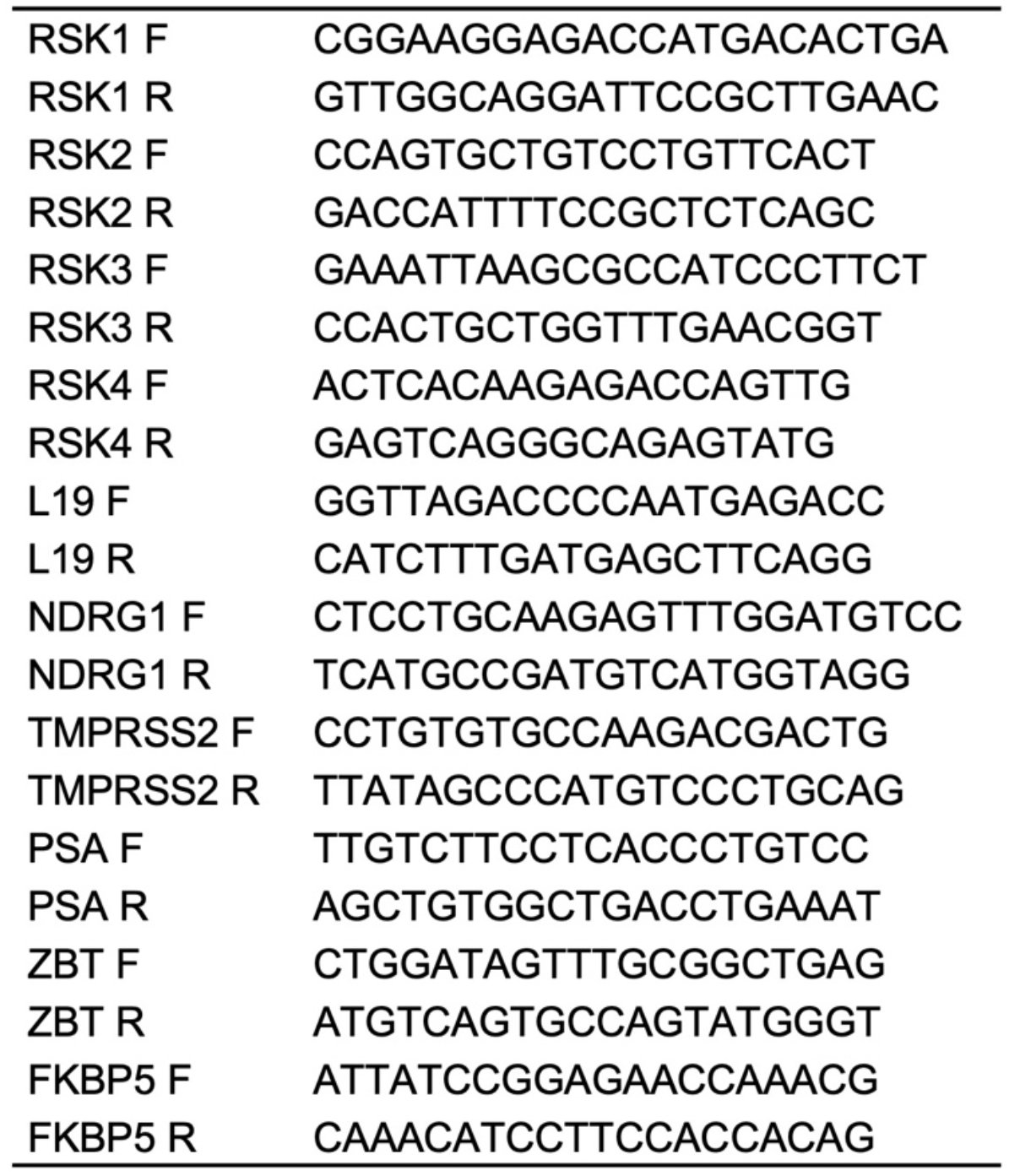
qPCR primers.

**Figure S1.**
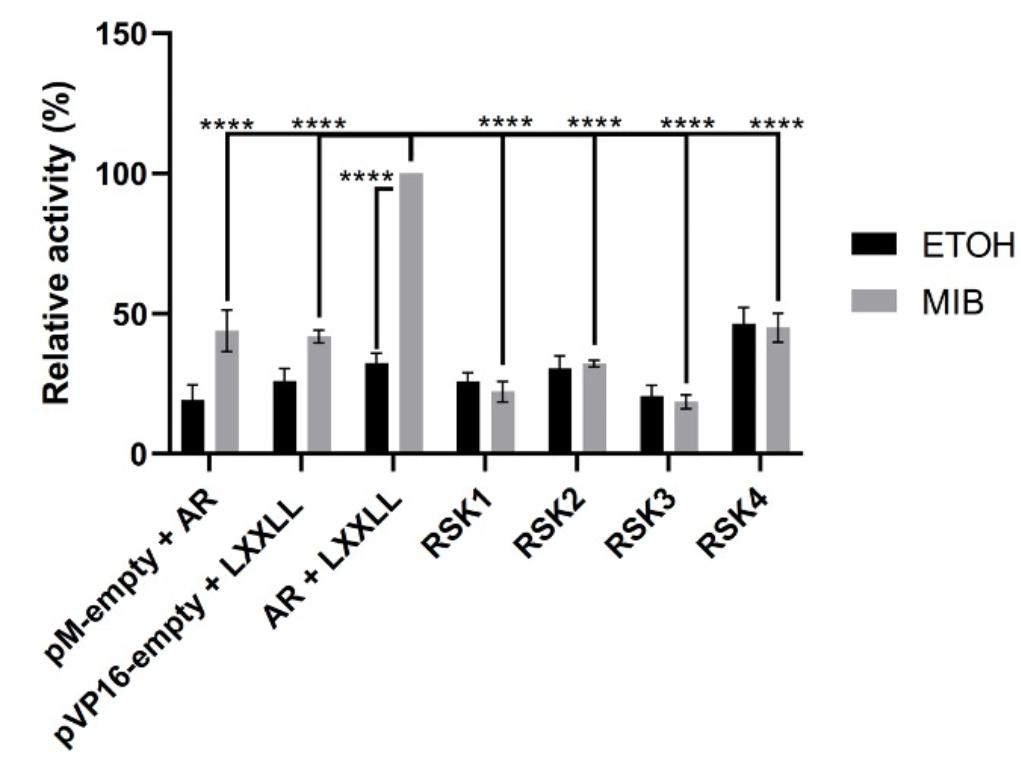
Mammalian 2-hybrid positive and negative controls. COS-1 cells were transfected with pVP16-AR, pM-LxxLL, pM-RSKs or empty plasmids. Luciferase activity was normalised to β-galactosidase activity and made relative to pVP16-AR + pM-LXXLL in the presence of MIB. Cells were treated with vehicle control (ethanol, ETOH, grey bar) or mibolerone (MIB, black bar). Data are presented as means ± standard deviation (SD). ANOVA comparison test was used to analyse the differences among multiple groups.^****^P < 0.0001.

**Table S2:** RSK3 and RSK4 fold increase in PCa cell lines vs PN1TA.

|  | RSK3 | RSK4 |
| --- | --- | --- |
| LNCAP | 40.9 | 4.9 |
| C42 | 24.6 | 6.9 |
| C42B | 15.9 | 5.2 |
| 22RV1 | NS | 9.1 |
| DU145 | 16.0 | NS |

NS = non significant

**Figure S2.**
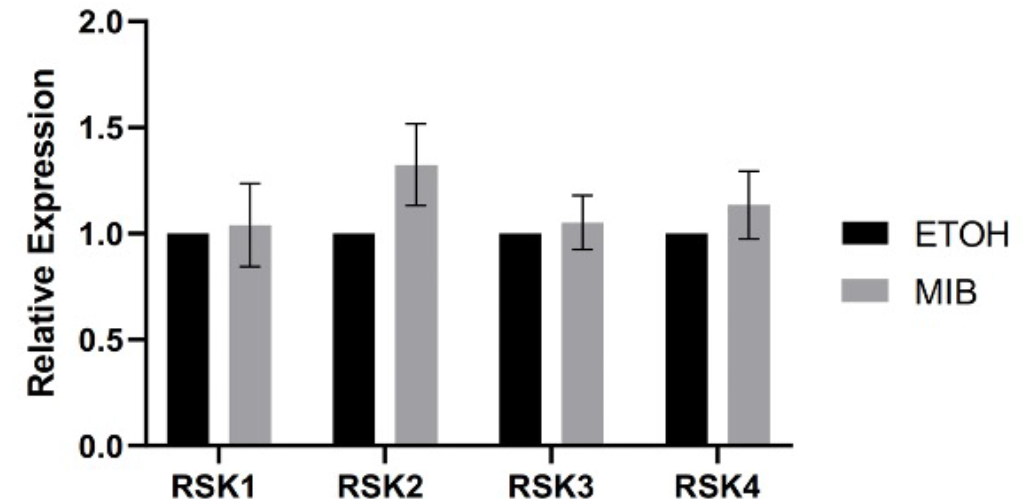
Androgen receptor signalling does not alter the expression of the RSKs. LNCaP cells were treated with vehicle control (ethanol, ETOH, grey bar) or mibolerone (MIB, black bar). qPCR analysis of total RNA extracted from cells was used to quantify RSK1/2/3/4. Data are the means from 3 independent biological repeats. Data are presented as means ± standard deviation (SD). ANOVA comparison test was used to analyse the differences among multiple groups, none showing statistically significant differences.

**Figure S3.**
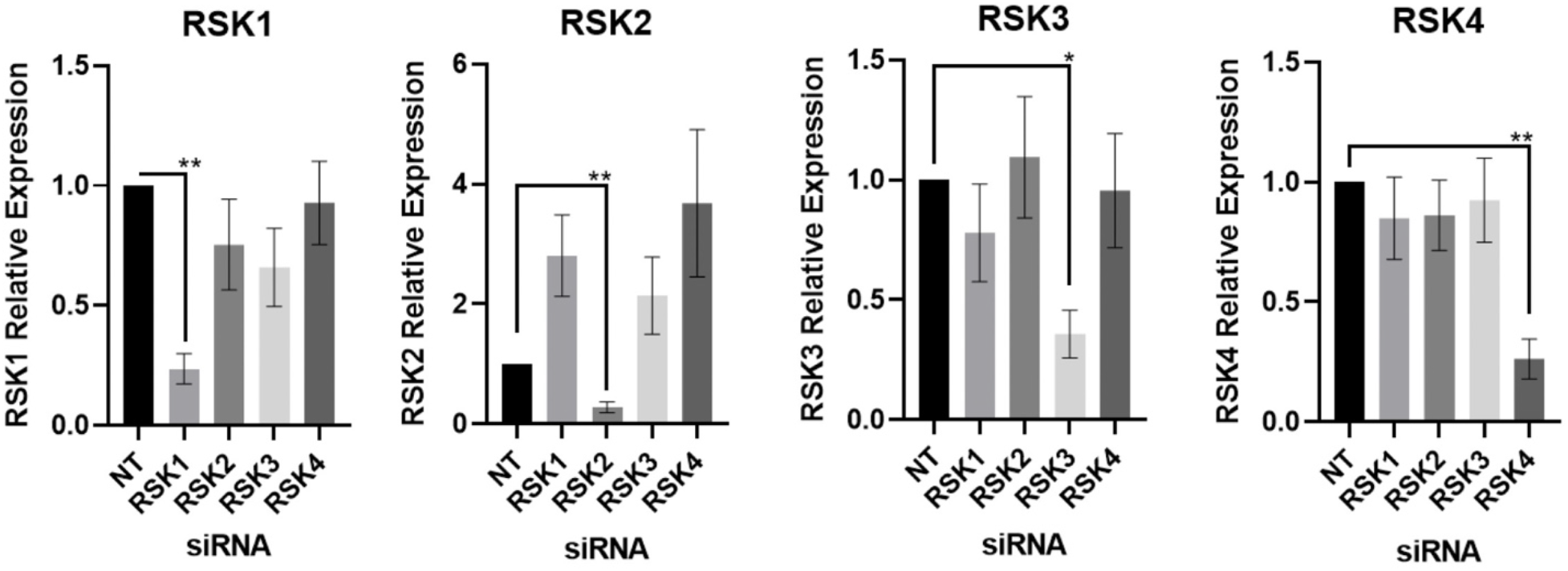
Confirmation of successful knockdown of the RSKs using siRNA. LNCaP cells were transfected with RSK1/2/3/4 and NT siRNAs. qPCR analysis of total RNA extracted from cells was used to quantify RSK1/2/3/4 expression. Data are the means from 3 independent biological repeats ± standard deviation (SD). ANOVA comparison test was used to analyse the differences among multiple groups. ^*^P < 0.05, ^**^P < 0.01.

**Figure S4.**
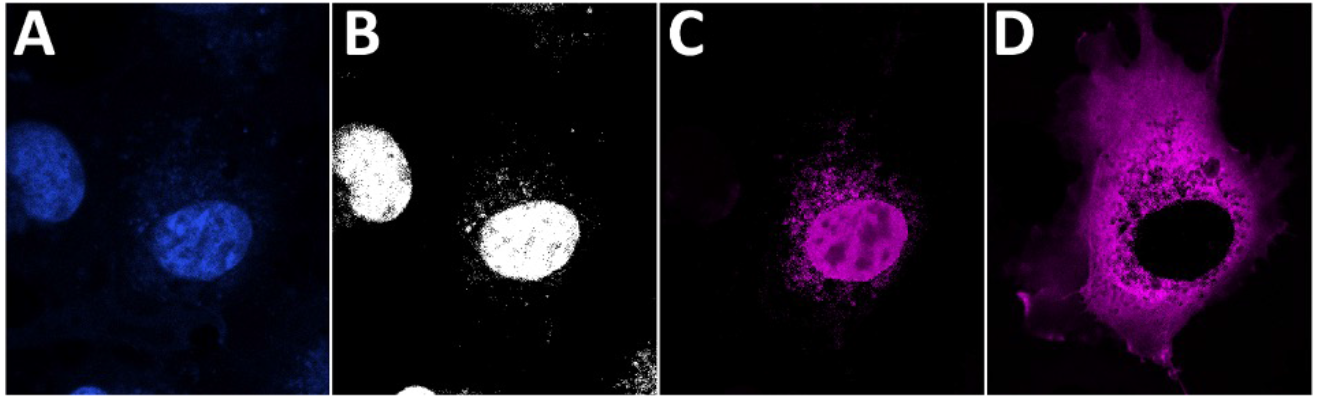
Image processing to determine nucleus/cytosol ratio. **A**), blue channel revealing the nucleus stained with the DNA-intercalating dye DAPI. **B**) Nucleus selection stored in the region of interest (ROI) manager. **C**) fluorescence intensity signal within the nucleus of the magenta channel showing mCherry-RSK1/2/3/4 labelled cells. **D**) Fluorescence intensity signal across the cytosol in the magenta channel.

**Table S3A.** p-values of the RSKs relative differences in the nucleus/cytosol ratio when co-expressed with AR.

|  | Condition | difference | CI_lo | CI_hi | p_median |
| --- | --- | --- | --- | --- | --- |
| 1 | RSK1 AR ETOH |  |  |  | 1 |
| 2 | RSK1 AR MIB | -0.02 | -0.188 | 0.122 | 0.901 |
| 3 | RSK2 AR ETOH | -0.046 | -0.243 | 0.138 | 0.75 |
| 4 | RSK2 AR MIB | 0.356 | 0.114 | 0.594 | 0.004 |
| 5 | RSK3 AR ETOH | 0.094 | -0.065 | 0.247 | 0.158 |
| 6 | RSK3 AR MIB | 0.123 | -0.044 | 0.296 | 0.108 |
| 7 | RSK4 AR ETOH | 0.141 | -0.024 | 0.29 | 0.079 |
| 8 | RSK4 AR MIB | 0.199 | 0.027 | 0.482 | 0.104 |

**Table S3B.**
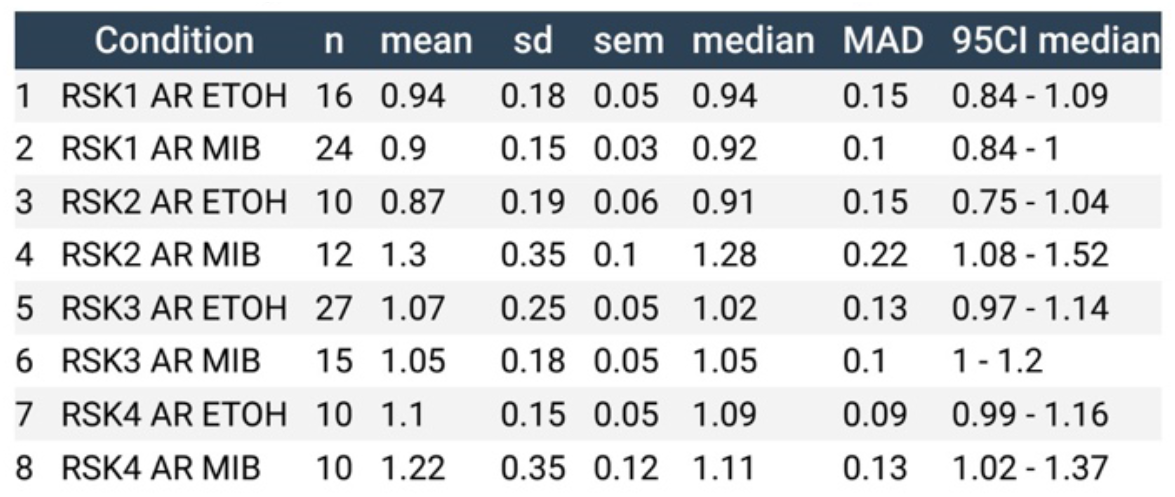
95% CL of the RSKs relative differences in the nucleus/cytosol ratio when co-expressed with AR.

|  | Condition | n | mean | sd | sem | median | MAD | 95CI median |
| --- | --- | --- | --- | --- | --- | --- | --- | --- |
| 1 | RSK1 AR ETOH | 16 | 0.94 | 0.18 | 0.05 | 0.94 | 0.15 | 0.84 - 1.09 |
| 2 | RSK1 AR MIB | 24 | 0.9 | 0.15 | 0.03 | 0.92 | 0.1 | 0.84 - 1 |
| 3 | RSK2 AR ETOH | 10 | 0.87 | 0.19 | 0.06 | 0.91 | 0.15 | 0.75 - 1.04 |
| 4 | RSK2 AR MIB | 12 | 1.3 | 0.35 | 0.1 | 1.28 | 0.22 | 1.08 - 1.52 |
| 5 | RSK3 AR ETOH | 27 | 1.07 | 0.25 | 0.05 | 1.02 | 0.13 | 0.97 - 1.14 |
| 6 | RSK3 AR MIB | 15 | 1.05 | 0.18 | 0.05 | 1.05 | 0.1 | 1 - 1.2 |
| 7 | RSK4 AR ETOH | 10 | 1.1 | 0.15 | 0.05 | 1.09 | 0.09 | 0.99 - 1.16 |
| 8 | RSK4 AR MIB | 10 | 1.22 | 0.35 | 0.12 | 1.11 | 0.13 | 1.02 - 1.37 |

**Table S3C.** p-values of the RSKs relative differences in the nucleus/cytosol ratio.

|  | Condition | difference | CI_lo | CI_hi | p_median |
| --- | --- | --- | --- | --- | --- |
| 1 | RSK1 |  |  |  | 1 |
| 2 | RSK2 | 0.008 | -0.232 | 0.346 | 0.998 |
| 3 | RSK3 | -0.226 | -0.459 | -0.011 | 0.027 |
| 4 | RSK4 | -0.092 | -0.308 | 0.137 | 0.392 |

**Table S3D.** 95% CL of the RSKs relative differences in the nucleus/cytosol.

|  | Condition | n | mean | sd | sem | median | MAD | 95CI median |
| --- | --- | --- | --- | --- | --- | --- | --- | --- |
| 1 | RSK1 | 10 | 1.07 | 0.18 | 0.06 | 1.07 | 0.16 | 0.92 - 1.26 |
| 2 | RSK2 | 10 | 1.13 | 0.33 | 0.11 | 1.06 | 0.2 | 0.93 - 1.33 |
| 3 | RSK3 | 10 | 0.85 | 0.15 | 0.05 | 0.86 | 0.14 | 0.71 - 0.95 |
| 4 | RSK4 | 10 | 1.01 | 0.18 | 0.06 | 0.96 | 0.1 | 0.87 - 1.19 |

**Figure S5.**
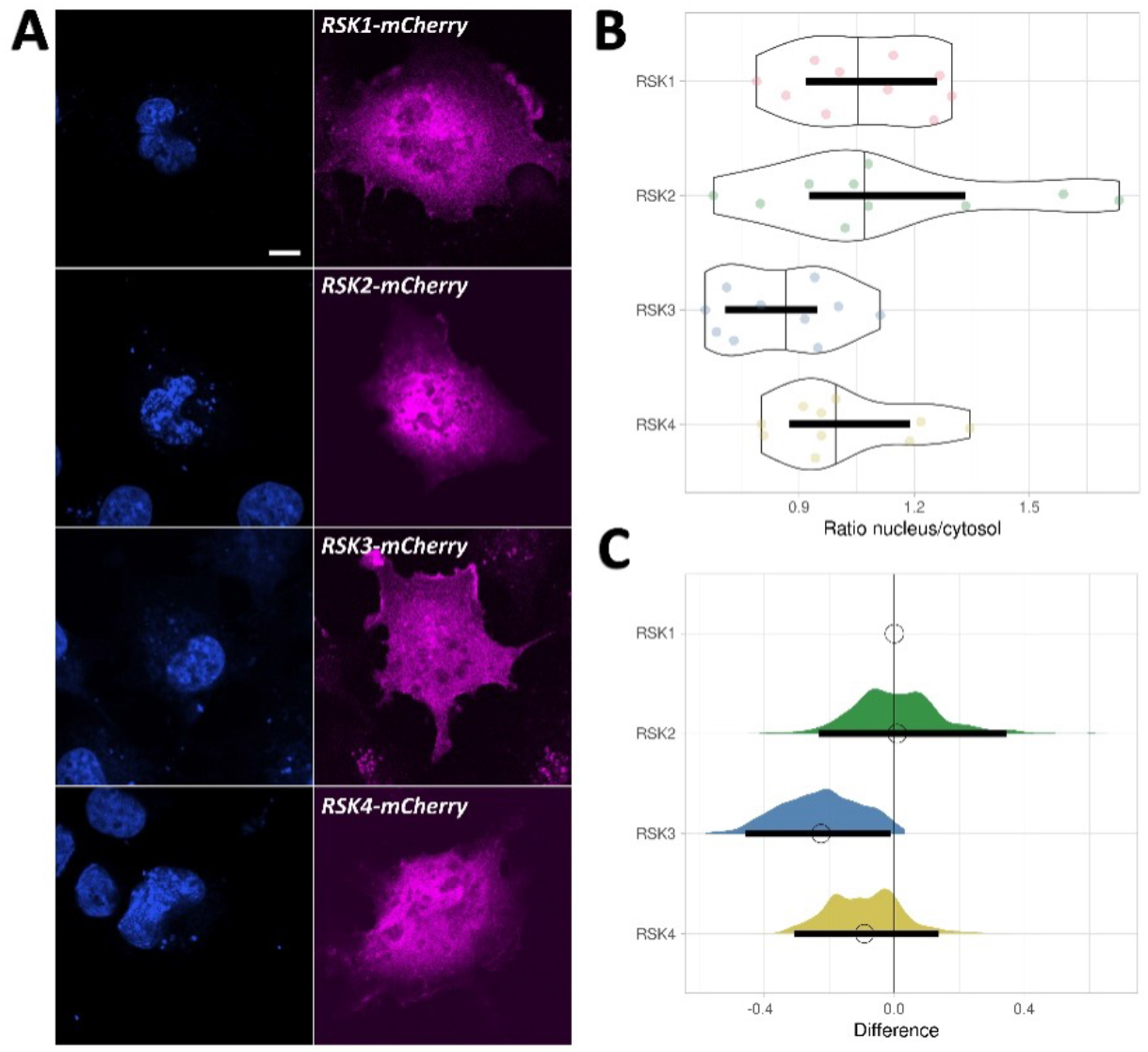
RSKs cellular localisation. **A)** Expression of mCherry-RSK1/2/3/4 constructs. Colour channels are arranged in columns, with DAPI-stained nuclei (blue), eGFP-AR (green), and, from top to bottom, mCherry-RSK1/2/3/4 (magenta). Scale bar indicates 10 μm. **B**) Violin plots of nucleus/cytosol ratio of mean fluorescence intensity for each of the RSKs (RSK1/2/3/4 from top to bottom). Thick black lines show the 95% compatibility interval (CI). The data points used to produce the box plots are overlaid in different colour. **C**) Relative differences in the nucleus/cytosol ratio of mean fluorescence intensity for RSK1/2/3/4 (same colours as in (B)), shown as relative effect sizes. The difference between mean values is determined relative to mCherry-RSK1, indicated with a circle at the top. The magenta double arrow on the top of the panel indicates a stronger cytosolic localisation on the left and nuclear on the right. The 95% CI is derived from the bootstrap distribution and indicated with the black vertical bars. See Table S3C-D for p-values and 95% CI.

**Figure S6.**
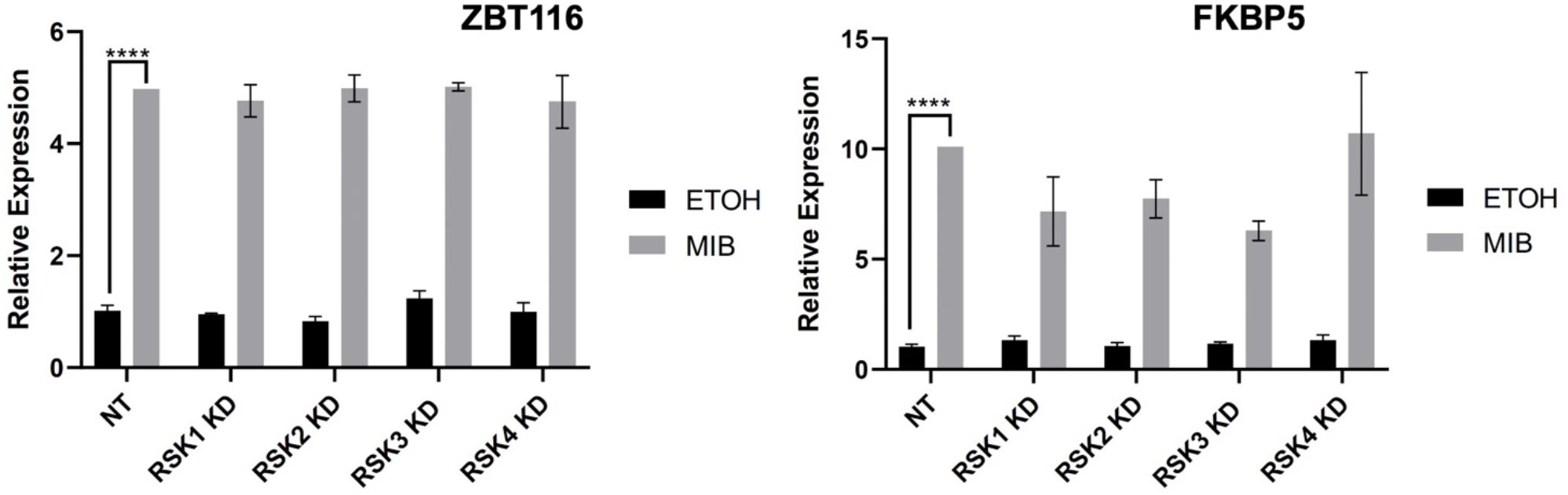
RSKs knock-down does not affect the expression of *ZBT116* and *FKBP5*. LNCaP cells were transfected with RSK1/2/3/4 and NT siRNAs, and treated with vehicle control (ethanol, ETOH, grey bar) or mibolerone (MIB, black bar). Q-RT-PCR analysis of total RNA Table S1A: cloning primers extracted from cells was used to quantify AR target genes expression, *ZBT116* and *FKBP5*. Table S1B: qPCR primers Data are the means from 3 independent biological repeats. Data are presented as means ± standard deviation (SD). ANOVA comparison test was used to analyse the differences among multiple groups. ^****^P < 0.0001.

## Notes

### Competing Interest Statement

The authors have declared no competing interest.

